# The Subtype Specificity of Genetic Loci Associated with Stroke in 16,664 cases and 32,792 controls

**DOI:** 10.1101/371153

**Authors:** Matthew Traylor, Christopher D. Anderson, Loes C.A. Rutten-Jacobs, Guido J. Falcone, Mary E. Comeau, Hakan Ay, Cathie L.M. Sudlow, Huichun Xu, Braxton D. Mitchell, John Cole, Kathryn Rexrode, Jordi Jimenez-Conde, Reinhold Schmidt, Raji P. Grewal, Ralph Sacco, Marta Ribases, Tatjana Rundek, Jonathan Rosand, Martin Dichgans, Jin-Moo Lee, Carl D. Langefeld, Steven J. Kittner, Hugh S. Markus, Daniel Woo, Rainer Malik, on behalf of the NINDS Stroke Genetics Network (SiGN) and International Stroke Genetics Consortium (ISGC)

## Abstract

**Background:** Genome-wide association studies have identified multiple loci associated with stroke. However, the specific stroke subtypes affected, and whether loci influence both ischaemic and haemorrhagic stroke, remains unknown. For loci associated with stroke, we aimed to infer the combination of stroke subtypes likely to be affected, and in doing so assess the extent to which such loci have homogeneous effects across stroke subtypes.

**Methods:** We performed Bayesian multinomial regression in 16,664 stroke cases and 32,792 controls of European ancestry to determine the most likely combination of stroke subtypes affected for loci with published genome-wide stroke associations, using model selection. Cases were subtyped under two commonly used stroke classification systems, Trial of Org 10172 Acute Stroke Treatment (TOAST) and Causative Classification of Stroke (CCS). All individuals had genotypes imputed to the Haplotype Reference Consortium 1.1 Panel.

**Results:** Sixteen loci were considered for analysis. Seven loci influenced both haemorrhagic and ischaemic stroke, three of which influenced ischaemic and haemorrhagic subtypes under both TOAST and CCS. Under CCS, 4 loci influenced both small vessel stroke and intracerebral haemorrhage. An *EDNRA* locus demonstrated opposing effects on ischaemic and haemorrhagic stroke. No loci were predicted to influence all stroke subtypes in the same direction and only one locus (12q24) was predicted to influence all ischaemic stroke subtypes.

**Conclusions:** Heterogeneity in the influence of stroke-associated loci on stroke subtypes is pervasive, reflecting differing causal pathways. However, overlap exists between haemorrhagic and ischaemic stroke, which may reflect shared pathobiology predisposing to small vessel arteriopathy. Stroke is a complex, heterogeneous disorder requiring tailored analytic strategies to decipher genetic mechanisms.

## Introduction

The burden of stroke on global healthcare and society is substantial; it is consistently one of the leading causes of death and disability worldwide, [1] and a major cause of cognitive impairment and dementia. However, there exist significant gaps in our understanding of the pathological processes that underlie the disease. In recent years genome-wide association studies (GWAS) have made considerable advances in identifying genetic components underlying complex traits, in many cases identifying novel disease pathways and treatments.[2]

Characterizing the genetic component to stroke has been challenging, in part due to clinical heterogeneity, with at least three distinct major pathological processes (cardioembolism, large artery atherosclerosis, small vessel disease) underlying the majority of ischaemic strokes; and two processes underlying the majority of intracerebral haemorrhagic stroke (small vessel disease and cerebral amyloid angiopathy). [3, 4] However, recent GWAS have made considerable advances; 32 independent genome-wide significant loci were identified in the MEGASTROKE project. [5] The majority of these loci were identified as being associated with inclusive ‘all stroke’ or ‘ischaemic stroke’ categories, rather than specific stroke subtypes. This is in part due to study design, with much larger samples for these broader categories and only a fraction of stroke cases having detailed phenotyping. Indeed, this finding is in contrast to earlier studies that identified loci such as *HDAC9, PITX2* as being associated with specific subtypes. [6, 7] In order to interpret genetic risk associations in the context of biological mechanisms, a pertinent question is whether the newly identified stroke-associated loci truly confer risk across all stroke subtypes, or whether isolated or combinations of subtypes are affected. At least one of the novel variants (on chromosome 1q22) shows association with both ischaemic and haemorrhagic stroke, which might point to some shared mechanisms underlying these clinically distinct entities, which have thus far been separated in genetic studies.

Conventional approaches to GWAS, which employ within study analysis and subsequent meta-analysis across groups, do not enable detailed model comparison across different subgroups. In this analysis, we used multinomial logistic regression on well-characterized subjects with individual-level data to investigate the association of all identified genetic GWAS loci to date with all stroke subtypes (cardioembolic (CES), large artery stroke (LAS), small vessel stroke (SVS) and intracerebral haemorrhage (ICH)), determining the most likely combination of stroke subtypes affected at each locus. We performed our analysis using two established subtyping approaches: the Trial of Org 10172 in Acute Stroke Treatment (TOAST), [8] and Causative Classification of Stroke (CCS) system,[9] to provide a comprehensive account of these loci across available classification systems. Our overall aim was to evaluate genetic loci identified in previous studies using stroke datasets with well-defined phenotyping to determine if subtype specificity or cross-subtype associations could be identified.

## Methods

### Cohort Characteristics

The data used in this analysis were derived from several sources: the NINDS-SIGN Stroke Genetics study, [10] the Wellcome Trust Case Control Consortium 2 Stroke and Immunochip studies, [6, 11] the UK Young Lacunar Stroke Study, [12] Genetics of Cerebral Hemorrhage with Anticoagulation (GOCHA), [13] Genetic and Environmental Risk Factors for Hemorrhagic Stroke (GERFHS), [13] Cambridge ICH Genetics Study. Almost all samples (>95%) were included in the previous MEGASTROKE genome-wide association study of stroke. [5]

### Stroke Phenotyping

Stroke was defined according to the World Health Organization (WHO), i.e. rapidly developing signs of focal (or global) disturbance of cerebral function, lasting more than 24 hours or leading to death with no apparent cause other than that of vascular origin. Strokes were defined as ischaemic stroke (IS) or intracerebral haemorrhage (ICH) based on clinical and imaging criteria. ICH stroke events were divided into lobar or deep, which have different presumed etiology, [3] based on location of the primary event. Ischaemic stroke cases were classified under the TOAST or CCS stroke classification systems (causative and phenotypic), or both. [8, 9] TOAST and CCS both include an ‘undetermined ischaemic stroke’ group (UND) denoting individuals for which it is not possible to determine the ischaemic stroke subtype. Full details are provided in Additional Files 1-2.

### Genotyping and Imputation

Genotyping of datasets has been described in detail elsewhere. [6, 10-13] In this analysis, we imputed all datasets to the Haplotype Reference Consortium 1.1 panel, using the Michigan Imputation Server. [14] For each separately imputed dataset, we extracted SNPs with MAF>1% and imputation INFO values>0.8. All datasets were subsequently merged using bcftools and SNPs with a MAF>5% in the combined dataset and present in 66% of samples were included in further analyses.[15] We removed any duplicate or related (3^rd^ degree or closer) samples at this stage and calculated ancestry informative principal components on a linkage-disequilibrium pruned subset of SNPs on the remaining individuals using the --pca approx function in plink 2.0.[16]

### Locus and SNP Selection

For each locus associated with stroke or stroke subtypes at genome-wide significance in MEGASTROKE,[5] we identified all SNPs in LD (r^2^>0.2) with the lead reported SNP based on the five European populations from 1000 Genomes.[17] These SNPs were then extracted from the merged dataset for analysis. We did not analyse two regions from MEGASTROKE: *RGS7* and *TMFSF1-TMFSF4*, as the previously associated variants in these regions were low frequency variants that were filtered out in our analysis. We additionally considered the COL4A2 locus as it been robustly associated with stroke phenotypes in other large-scale studies. [18]

### Multinomial Logistic Regression

We used a Bayesian multinomial logistic regression approach, implemented in *Trinculo*, [19] to evaluate the association of SNPs at each locus. Multinomial logistic regression is a natural extension of logistic regression that enables modelling of multiple phenotypic categories simultaneously against a common set of controls. The benefit of this approach, which is leveraged in this analysis, is that it enables comparison of models that include different combinations of phenotypes. In the context of genetic studies, this enables determination of the combination of phenotypes that are mostly likely to be associated with the genetic variant of interest.

We used the default prior, which assumes effect sizes are independent with variances of 0.04. All analyses included eight ancestry-informative principal components, and batch covariates for each study.

Based on their association at genome-wide significance in previous analyses, we assumed a *priori* that each region was associated with stroke. However, to avoid overfitting for weakly associated loci in our data, we performed model selection only for loci that had a Bayes Factor of at least 4 in either TOAST or CCS analyses.

No prior genome-wide association study of stroke has identified a significant association with strokes of undetermined or cryptogenic cause. Given that this study was intended to evaluate potential shared mechanisms between subtypes, we excluded strokes of undetermined cause in model fitting.

### Statistical Analysis

For each locus we performed the following steps:

1. Use multinomial logistic regression to model the association between each genetic variant and stroke subtypes under TOAST and CCS classifications, in each case including ICH as an additional outcome. We therefore tested a common set of Controls against CES, LAS, SVS, UND, and ICH cases.
2. Identify the most significant SNP in the locus under any classification system
3. For this SNP, calculate marginal likelihoods for all combinations of phenotypes
4. Identify the combination of phenotypes with the largest marginal likelihood (discarding any groups containing UND) and infer that this indicates the most likely combination of phenotypes for which the SNP confers risk

## Results

After QC, there were up to 16,664 cases and 32,792 controls remaining for analysis (Table 1). In the merged dataset, a binomial genome-wide analysis of all cases against controls had a genomic inflation lambda=1.09, while the LDSCORE intercept value was 1.04, [20] suggesting that the majority of inflation was due to polygenicity and that any bias introduced by merging the datasets was minimal. A comparison of odds ratios for analysed loci from MEGASTROKE and the most recent ICH publication with those from our analysis showed high consistency (r^2^=0.95, Additional File 3) despite slightly differing samples.

**Table 1.** Sample Sizes

| Classification System | CES | LAS | SVS | UND | ICH | Controls |
| --- | --- | --- | --- | --- | --- | --- |
| TOAST | 3847 | 2803 | 3976 | 4085 | 1953 | 32,792 |
| CCS | 2826 | 2204 | 3093 | 4013 | 1953 | 28,052 |

CES, cardioembolic Stroke; LAS, large artery atherosclerotic stroke; SVS, small artery occlusion stroke; UND, stroke of undetermined etiology; ICH, intracerebral haemorrhage; TOAST, Trial of Org 10172 Acute Stroke Treatment Classification System; CCS, Causative Classification of Stroke System (causative system).

Sixteen loci contained SNPs with Bayes factors of at least 4 in either TOAST or CCS analyses. We took these sixteen loci forward for further model selection. Plots for all loci under each classification system are provided in Additional Files 4-19. For each of the sixteen loci, we identified the most likely combination of associated phenotypes at each locus (Figure 1) based on model selection. Apart from one locus (*FOXF2*), we found identical results between the two CCS systems, so for simplicity of presentation results for CCS causative are presented only.

**Figure 1.**
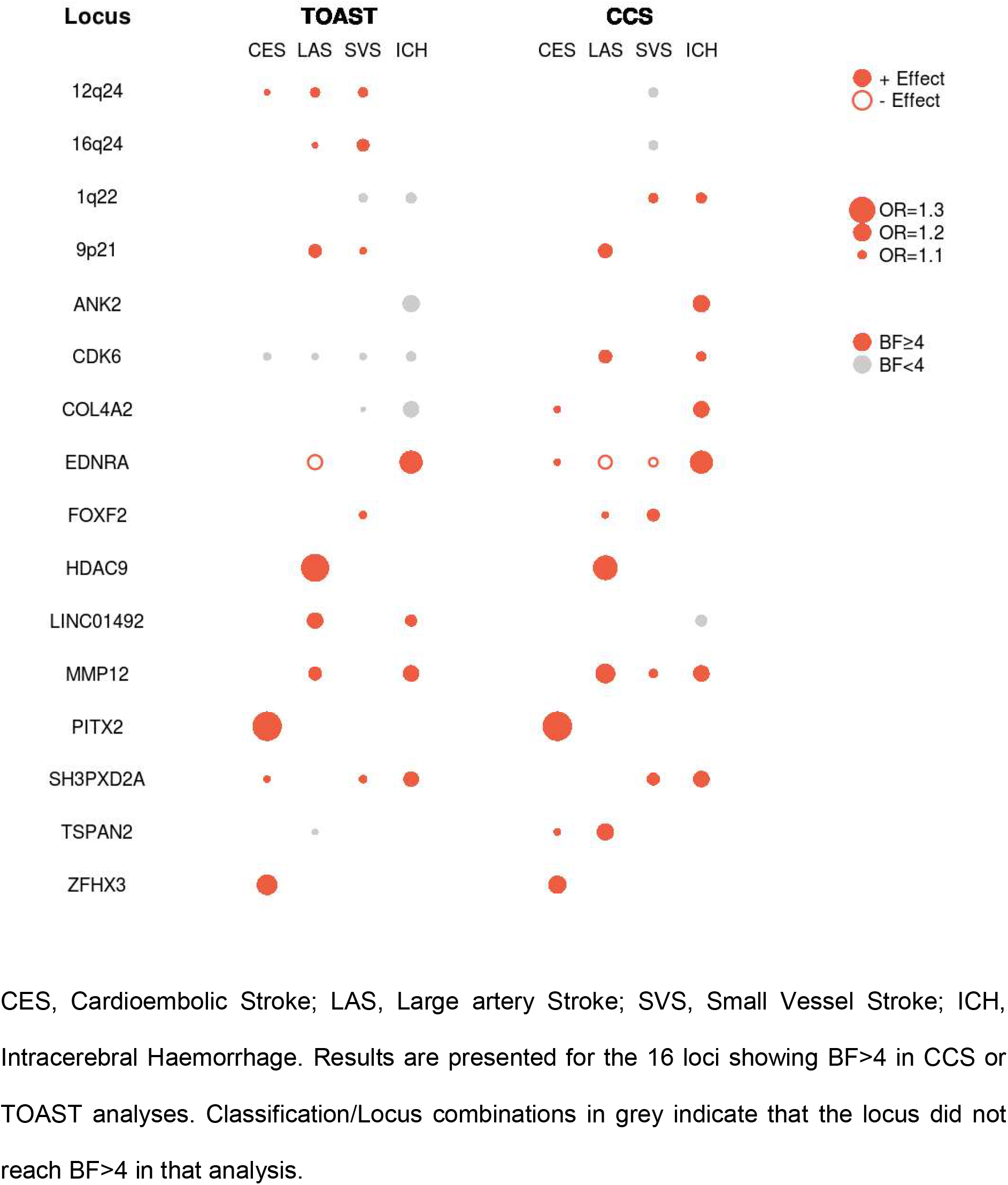
Stroke Subtypes in Best Fitting Model at Each Locus, for CCSc, CCSp, and TOAST classification Systems, with Size Weighted by Association Odds Ratio

For seven loci, the combination of phenotypes most likely to be influenced by the lead genetic variant at the loci included both ischaemic and haemorrhagic stroke subtypes. Four of these are shown in Figure 2. At these four loci*: EDNRA, 1q22, MMP12, SH3PXD2A*, the ischaemic subtype included SVS, highlighting shared mechanisms underlying ICH and SVS, likely through predisposition to cerebral small vessel disease. At the *EDNRA* locus, the direction of association for ICH was opposite to that for LAS and SVS, pointing to contrasting influence on ischaemic and haemorrhagic stroke risk. We explored whether ICH-associated loci were specific to deep or lobar ICH. As in previous reports, [13, 18] associations at 1q22 and COL4A2 appear to be specific to deep ICH, with no effect in lobar ICH. For other regions, the evidence for specificity was more equivocal (Additional File 20).

**Figure 2.**
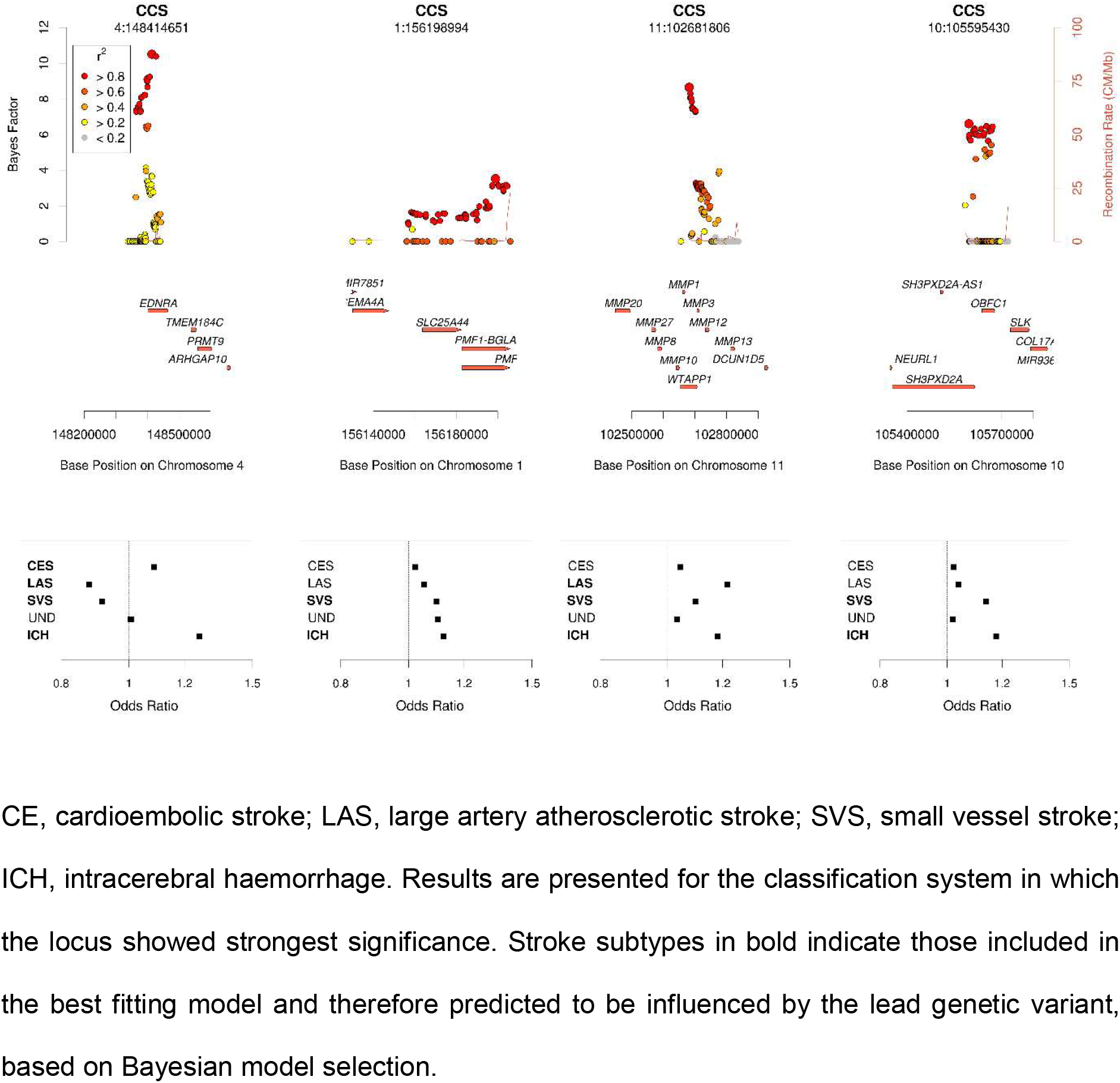
Local Plots showing Associations with 4 Regions Conferring Risk of Ischaemic and Haemorrhagic Stroke and Odds Ratios for all stroke Subtypes

For four loci: *HDAC9, PITX2, ZFHX3, ANK2*, only one phenotype was affected by the lead variant (Figure 1, Additional Files 13, 16, 19, 8) in the most likely configuration across all classification systems. Several other loci: 9p21, 12q24, 16q24, FOXF2 were associated with only one phenotype under particular classification systems, but did not show consistency across TOAST and CCS (Additional Files 5, 6, 7, 12). For *TSPAN2*, which was previously identified as being associated with LAS, [10] the best-fit model also included CES under CCS, albeit with a much weaker effect than LAS (rs17479660; CES, OR=1.08; LAS, OR=1.19 under CCS). Echoing previous results, the locus showed much stronger significance under CCS classifications than under TOAST (Additional File 18).

For *COL4A2*, the strongest association found under TOAST was for rs9515201. The most likely model contained ICH (OR=1.14) and SVS (OR=1.13), consistent with findings from previous analyses. [18] However, under CCS an alternate SNP, rs1927349, was the strongest associated. No association with SVS was observed, and a weak association with CES was observed instead. Reasons for this discrepancy between CCS and TOAST are not immediately clear, but non-overlapping samples between the two classification systems are a likely factor.

The mean (SD) number of stroke subtypes affected at each locus were 1.88 (0.89) under TOAST and 1.69 (0.87) under CCS. Under CCS, the most common combination of affected subtypes was SVS and ICH (4 loci).

## Discussion

We performed a large-scale genetic analysis, characterising the effects of established stroke risk loci with ischaemic and haemorrhagic stroke subtypes in up to 16,664 cases and 32,792 controls.

Our main findings are twofold. First, for the vast majority of loci studied, multiple but never all stroke subtypes were affected at the locus. Only one locus (12q24) was assumed to influence all ischaemic stroke subtypes. This indicates that although these loci were identified in analyses of inclusive stroke phenotypes, in the main their effects are specific to particular combinations of stroke subtypes. The mean number of subtypes affected was 1.88 for TOAST and 1.69 for CCS classification systems. Notable exceptions were the *PITX2* and *ZFHX3* loci, which were associated with cardioembolic stroke most likely through atrial fibrillation, and *HDAC9* which is associated with large vessel stroke. Under TOAST, the *FOXF2* locus was associated solely with SVS. However, under CCS, LAS was also implicated. For CCS, the 9p21 locus was predicted to influence only LAS. However, under TOAST, SVS was also implicated. Our analyses suggest that *ANK2* confers risk of stroke predominantly through its influence on *ICH.* We were unable to identify any loci for which the most likely model included all stroke phenotypes in the same direction and only one (12q24) which for which the most likely model included all ischaemic stroke subtypes.

Secondly, we find evidence that several loci influence both haemorrhagic and ischaemic stroke. This was evident for seven loci in total (1q22, *COL4A2*, *EDNRA*, *LINC01492*, *MMP12*, *SH3PXD2A*, *CDK6*). Under CCS, 4 loci (*SH3PXD2A*, *MMP12*, *EDNRA*, 1q22) influenced both SVS and ICH, highlighting shared mechanisms underlying small vessel disease. Previous GWAS analyses have tended to separate ischaemic and haemorrhagic stroke on the basis of presumed differing etiologies. Our results suggest that including haemorrhagic alongside ischaemic stroke in multiphenotype analyses will provide further insights.

For one locus: Endothelin Receptor Type A (*EDNRA*), the association with ICH was in the opposite direction to the ischaemic stroke subtypes, suggesting opposing risk mechanisms. This locus has previously been associated with a variety of vascular phenotypes, including coronary artery disease, carotid plaques, and peripheral arterial disease (in concordant direction with ischaemic stroke), as well as intracranial aneurysm (in concordant direction with intracerebral haemorrhage). [21-24] The locus has also been associated with migraine in candidate gene studies, [25] but this has not been validated in GWA studies. [26] EDNRA encodes the type A receptor (*ET_A_*) for Endothelin-1 (*ET-1*), a potent vasoconstrictor with pro-inflammatory effects. *ET_A_* -specific antagonists increase Nitric Oxide (NO)-mediated endothelium-dependent relaxation, reduce ET-1 levels and inhibit atherosclerosis in mice, [27] suggesting that higher levels of *ET_A_* are pro-atherogenic: consistent with the observation that higher *ET_A_* levels are observed in atherosclerotic plaques. [28] This is also consistent with the C allele of rs17612742 in our study leading to increased risk of ischaemic stroke through elevated *ET_A_* levels. Indeed, in GWA studies of intracranial aneurysm the susceptibility variant (in LD with the T allele of rs17612742 in our study) was shown to result in higher transcription factor binding affinity, likely resulting in repression of the transcriptional activity of *EDNRA*. [23] The reason why lower levels of *ET_A_* might promote intracranial aneurysm and intracerebral haemorrhage is not immediately obvious, but several mechanisms are possible. Levels of *ET-1* have been linked to vascular remodelling, an important process underlying ICH and IA; [29, 30] subtle changes in this process induced by altered availability of *ET_A_* is one such mechanism. Deep ICH and ischaemic SVS arise due to the same arteriopathy that arises in the deep perforating arteries of the brain. The *EDNRA* variant in this study points to a mechanism that influences whether the resulting pathology is ischaemic or haemorrhagic, and as such warrants further detailed investigation.

Some loci were notably more significant when phenotyped using CCS; *SH3XPD2A, MMP12, TSPAN2, FOXF2, EDNRA*, which might point to CCS having greater accuracy and therefore utility in stroke GWA studies. However, the opposite was also true for others: *16q24, HDAC9*. We note that some differences may be due to the fact that not all individuals were subtyped under both CCS and TOAST; the TOAST cohort was a least 20% larger. A detailed discussion of the relative merits of TOAST and CCS is beyond the scope of this article, but our results highlight that the importance of collecting individual phenotypic qualities that make up the etiologic subtypes in genetic studies of stroke so that associated loci can be more systematically examined.

Our study has several strengths. The dataset was a large stroke population including intracerebral haemorrhage and ischaemic stroke cases, the majority of which were subtyped under both TOAST and CCS. We had full access to genotype-level data enabling us full control over all analyses. Similarly there are limitations. We present results for the most likely combination of stroke phenotypes affected at each locus: the ‘best-fitting’ model. We had limited statistical power to determine with statistical certainty that this was the correct model; significantly larger samples would be required to achieve this. Due to the challenges of performing these analyses across different ancestry populations, we performed analyses in European populations only. The results can therefore not be generalized to all populations. In all analyses we assume there is a single causal variant at the locus, which may not be true in all cases. Our analyses are based on use of a default prior, which has been used in many genetic studies. An alternative is to derive an empirical prior from associated genetic loci. As more loci are identified as being associated with stroke, this will become a more realistic possibility and should be explored in future analyses.

## Conclusions

Our findings suggest that although large scale genome-wide studies of broad ‘all stroke’ or ‘all ischaemic stroke’ phenotypes are able to identify multiple associations, it should not be assumed that such associations confer risk equally across stroke subtypes. Heterogeneity in the influence of genetic variants on different stroke subtypes is the norm, not the exception. Analyses such as the current one provide insights into the etiological stroke subtypes most prominently influenced by genetic variants at these loci – a prerequisite to decide on the most appropriate model systems to choose for further mechanistic studies. Stroke is a complex, heterogeneous disorder: our findings highlight the ongoing need for large, well phenotyped case collections and tailored analytic strategies to decipher the underlying genetic mechanisms.

## Abbreviations

CES: cardioembolic stroke
LAS: large artery stroke
SVS: small vessel stroke
ICH: intracerebral haemorrhage
SNP: single nucleotide polymorphism

## Acknowledgements

This work was supported by a British Heart Foundation Programme Grant (RG/16/4/32218). The NINDS-SIGN study was funded by the US National Institute of Neurological Disorders and Stroke, National Institutes of Health (U01 NS069208 and R01 NS100178). Collection of the UK Young Lacunar Stroke DNA Study (DNA Lacunar) was primarily supported by the Wellcome Trust (WT072952) with additional support from the Stroke Association (TSA 2010/01). Genotyping of the DNA Lacunar samples was supported by a Stroke Association Grant (TSA 2013/01). The principal funding for the WTCCC2 stroke study was provided by the Wellcome Trust, as part of the Wellcome Trust Case Control Consortium 2 project (085475/B/08/Z and 085475/Z/08/Z and WT084724MA). Hugh Markus is supported by a National Institute for Health Research (NIHR) Senior Investigator award, and his work is supported by the Cambridge Universities NIHR Comprehensive Biomedical Research Centre. Dr. Anderson is supported by NIH R01NS103924 and K23NS086873.

## Author’s Contributions

MT and RM designed the experiments. MT and MC performed the imputations. MT performed the statistical analyses. MT, CDA, LCARJ, HSM, DW, and RM wrote the first draft of the manuscript. All authors read and approved the final manuscript.

## Ethics approval and consent to participate

All research participants contributing clinical and genetic samples for analysis in this study provided written informed consent.

## Availability of data and materials

Data from the NINDS-SIGN Stroke study are available to researchers through dbGAP: https://www.ncbi.nlm.nih.gov/projects/gap/cgi-bin/study.cgi?study_id=phs000615.v1.p1. Trinculo v0.96 is available from: https://sourceforge.net/projects/trinculo/files/

## Additional Files

**Additional File 1.** Stroke Phenotyping

**Additional File 2.** Cohort Descriptions

**Additional File 3.** Comparison of log(odds ratio) from most recent publication with those from this analysis for 16 SNPs tested in this analysis

ICH, Intracerebral haemorrhage; CES, cardioembolic stroke; LAS, large artery stroke; SVS, small vessel stroke. Where the lead SNP from previous publication was not available, [5, 13] we used the nearest proxy (r^2^>0.6 in all cases). No SNPs in the 12q24 region passed QC in the most recent ICH publication so are not included here.

**Additional File 4.** 1q22 Region

**Additional File 5.** 9p21 Region

**Additional File 6.** 12q24 Region

**Additional File 7.** 16q24 Region

**Additional File 8.** ANK2 Region

**Additional File 9.** CDK6 Region

**Additional File 10.** COL4A2 Region

**Additional File 11.** EDNRA Region

**Additional File 12.** FOXF2 Region

**Additional File 13.** HDAC9 Region

**Additional File 14.** LINC01492 Region

**Additional File 15.** MMP12 Region

**Additional File 16.** PITX2 Region

**Additional File 17.** SH3PXD2A Region

**Additional File 18.** TSPAN2 Region

**Additional File 19.** ZFHX3 Region

**Additional File 20.** Odds ratios for association of ICH-associated loci with ICH subtypes, and evidence for ICH subtype-specific effects

OR, odds ratio; BF, Bayes Factor

## References

1. Global, regional, and national disability-adjusted life-years (DALYs) for 333 diseases and injuries and healthy life expectancy (HALE) for 195 countries and territories, 1990-2016: a systematic analysis for the Global Burden of Disease Study 2016. Lancet 2017, 390:1260–1344.

2. Visscher PM, Wray NR, Zhang Q, Sklar P, McCarthy MI, Brown MA, et al: 10 Years of GWAS Discovery: Biology, Function, and Translation. Am J Hum Genet 2017, 101:5–22.

3. Martini SR, Flaherty ML, Brown WM, Haverbusch M, Comeau ME, Sauerbeck LR, et al: Risk factors for intracerebral hemorrhage differ according to hemorrhage location. Neurology 2012, 79:2275–2282.

4. Hankey GJ: Stroke. Lancet 2017, 389:641–654.

5. Malik R, Chauhan G, Traylor M, Sargurupremraj M, Okada Y, Mishra A, et al: Multiancestry genome-wide association study of 520,000 subjects identifies 32 loci associated with stroke and stroke subtypes. Nat Genet 2018, 50:524–537.

6. Bellenguez C, Bevan S, Gschwendtner A, Spencer CC, Burgess AI, Pirinen M, et al: Genome-wide association study identifies a variant in HDAC9 associated with large vessel ischemic stroke. Nat Genet 2012, 44:328–333.

7. Traylor M, Farrall M, Holliday EG, Sudlow C, Hopewell JC, Cheng YC, et al: Genetic risk factors for ischaemic stroke and its subtypes (the METASTROKE collaboration): a meta-analysis of genome-wide association studies. Lancet Neurol 2012, 11:951–962.

8. Adams HP, Jr., Bendixen BH, Kappelle LJ, Biller J, Love BB, Gordon DL, et al: Classification of subtype of acute ischemic stroke. Definitions for use in a multicenter clinical trial. TOAST. Trial of Org 10172 in Acute Stroke Treatment. Stroke 1993, 24:35–41.

9. Ay H, Benner T, Arsava EM, Furie KL, Singhal AB, Jensen MB, et al: A computerized algorithm for etiologic classification of ischemic stroke: the Causative Classification of Stroke System. Stroke 2007, 38:2979–2984.

10. Loci associated with ischaemic stroke and its subtypes (SiGN): a genome-wide association study. Lancet Neurol 2016, 15:174–184.

11. Kilarski LL, Achterberg S, Devan WJ, Traylor M, Malik R, Lindgren A, et al: Meta-analysis in more than 17,900 cases of ischemic stroke reveals a novel association at 12q24.12. Neurology 2014, 83:678–685.

12. Traylor M, Bevan S, Baron JC, Hassan A, Lewis CM, Markus HS: Genetic Architecture of Lacunar Stroke. Stroke 2015, 46:2407–2412.

13. Woo D, Falcone GJ, Devan WJ, Brown WM, Biffi A, Howard TD, et al: Meta-Analysis of Genome-Wide Association Studies Identifies 1q22 as a Susceptibility Locus for Intracerebral Hemorrhage. Am J Hum Genet 2014, 94:511–521.

14. McCarthy S, Das S, Kretzschmar W, Delaneau O, Wood AR, Teumer A, et al: A reference panel of 64,976 haplotypes for genotype imputation. Nat Genet 2016, 48:1279–1283.

15. Danecek P, Auton A, Abecasis G, Albers CA, Banks E, DePristo MA, et al: The variant call format and VCFtools. Bioinformatics 2011, 27:2156–2158.

16. Chang CC, Chow CC, Tellier LC, Vattikuti S, Purcell SM, Lee JJ: Second-generation PLINK: rising to the challenge of larger and richer datasets. Gigascience 2015, 4:7.

17. Abecasis GR, Auton A, Brooks LD, DePristo MA, Durbin RM, Handsaker RE, et al: An integrated map of genetic variation from 1,092 human genomes. Nature 2012, 491:56–65.

18. Rannikmae K, Sivakumaran V, Millar H, Malik R, Anderson CD, Chong M, et al: COL4A2 is associated with lacunar ischemic stroke and deep ICH: Meta-analyses among 21,500 cases and 40,600 controls. Neurology 2017, 89:1829–1839.

19. Jostins L, McVean G: Trinculo: Bayesian and frequentist multinomial logistic regression for genome-wide association studies of multi-category phenotypes. Bioinformatics 2016, 32:1898–1900.

20. Bulik-Sullivan BK, Loh PR, Finucane HK, Ripke S, Yang J, Patterson N, et al: LD Score regression distinguishes confounding from polygenicity in genome-wide association studies. Nat Genet 2015, 47:291–295.

21. Bis JC, Kavousi M, Franceschini N, Isaacs A, Abecasis GR, Schminke U, et al: Meta-analysis of genome-wide association studies from the CHARGE consortium identifies common variants associated with carotid intima media thickness and plaque. Nat Genet 2011, 43:940–947.

22. Howson JMM, Zhao W, Barnes DR, Ho WK, Young R, Paul DS, et al: Fifteen new risk loci for coronary artery disease highlight arterial-wall-specific mechanisms. Nat Genet 2017, 49:1113–1119.

23. Low SK, Takahashi A, Cha PC, Zembutsu H, Kamatani N, Kubo M, et al: Genome-wide association study for intracranial aneurysm in the Japanese population identifies three candidate susceptible loci and a functional genetic variant at EDNRA. Hum Mol Genet 2012, 21:2102–2110.

24. Matsukura M, Ozaki K, Takahashi A, Onouchi Y, Morizono T, Komai H, et al: Genome-Wide Association Study of Peripheral Arterial Disease in a Japanese Population. PLoS One 2015, 10:e0139262.

25. Tzourio C, El Amrani M, Poirier O, Nicaud V, Bousser MG, Alperovitch A: Association between migraine and endothelin type A receptor (ETA-231 A/G) gene polymorphism. Neurology 2001, 56:1273–1277.

26. Gormley P, Anttila V, Winsvold BS, Palta P, Esko T, Pers TH, et al: Meta-analysis of 375,000 individuals identifies 38 susceptibility loci for migraine. Nat Genet 2016, 48:856–866.

27. Barton M, Haudenschild CC, d’Uscio LV, Shaw S, Munter K, Luscher TF: Endothelin ETA receptor blockade restores NO-mediated endothelial function and inhibits atherosclerosis in apolipoprotein E-deficient mice. Proc Natl Acad Sci U S A 1998, 95:14367–14372.

28. Ramzy D, Rao V, Tumiati LC, Xu N, Sheshgiri R, Miriuka S, et al: Elevated endothelin-1 levels impair nitric oxide homeostasis through a PKC-dependent pathway. Circulation 2006, 114:I319–326.

29. Penn DL, Witte SR, Komotar RJ, Sander Connolly E, Jr.: The role of vascular remodeling and inflammation in the pathogenesis of intracranial aneurysms. J Clin Neurosci 2014, 21:28–32.

30. Amiri F, Virdis A, Neves MF, Iglarz M, Seidah NG, Touyz RM, et al: Endothelium-restricted overexpression of human endothelin-1 causes vascular remodeling and endothelial dysfunction. Circulation 2004, 110:2233–2240.

